# Pathogenic potential in catheter-associated *Escherichia coli* is associated with separable biofilm and virulence gene determinants

**DOI:** 10.1101/2021.07.16.452752

**Authors:** Zongsen Zou, Robert F. Potter, William H. McCoy, George L. Katumba, Peter J. Mucha, Gautam Dantas, Jeffrey P. Henderson

## Abstract

Urinary catheterization facilitates *Escherichia coli* colonization of the urinary tract and increases infection risk. While specific pathotypes are well-recognized for some *E. coli* infections, it is unclear whether strain-specific characteristics among *E. coli* are associated with infection risk in catheterized patients. Here we used comparative genomics and a simulated catheter biofilm model to compare strains associated with catheter-associated urinary tract infection (CAUTI) and catheter-associated asymptomatic bacteriuria (CAASB). CAUTI was associated with a phylotype B2 sub-clade dominated by the multidrug resistant ST131 lineage, while CAASB isolates were genetically more diverse. Catheter-associated biofilm formation was widespread but quantitatively variable among isolates. Network community analysis resolved distinct groups of genes associated with infection or biofilm formation, with iron acquisition-associated genes prominent throughout. Using a reporter construct and targeted mutagenesis, we detected a biofilm phenotype for the ferric citrate transport (Fec) system, the most prominent correlate of high catheter biofilm formation in these patients. In mixed cultures, catheter biofilms formed by some CAASB strains suppressed catheter colonization by ST131 CAUTI isolates. These results are consistent with a paradigm in which catheter biofilm-associated genes increase infection risk in strains with a high pathogenic potential and decrease infection risk through niche exclusion in strains with low pathogenic potential.

## Introduction

Catheter-associated urinary tract infections (CAUTI) are among the most common nosocomial infections, with over one million cases annually in the United States [1–3]. Accurate diagnosis and effective treatment of CAUTI and ureteral stent-associated infections, can be challenging [4]. Bacteriuria alone is an insufficient criterion to establish a CAUTI diagnosis, which also requires attributable patient signs or symptoms such as suprapubic tenderness, flank pain, or fever [5, 6]. Patients with catheter-associated asymptomatic bacteriuria (CAASB) are at low risk of serious infection antibiotics are not recommended [7]. When symptoms are masked or of uncertain connection to the urinary tract, physicians must weigh the risk of progressive infection against the risks of catheter or device removal and inappropriate antibiotic therapy. In this context, professional society guidelines have long noted a need to better discriminate CAASB from CAUTI and to predict a patient’s risk for progression to CAUTI [8, 9].

The pathogenic potential of bacteria is a function of both host and bacterial characteristics [10]. In the urinary tract, the presence of a catheter or stent is an especially influential host characteristic, conferring a well-recognized predisposition to infection [2, 3]. By affecting urinary flow, providing an abiotic surface for bacterial adherence, and changing the local epithelium [11–13], these devices are associated with a distinctive physiology. The ability of *E. coli* to form biofilms on these devices is generally regarded as an important virulence characteristic in these patients [14, 15]. These biofilms are adherent bacterial communities enmeshed in an extracellular polymeric substance (EPS) matrix that form in response to specific environmental cues, permitting a resident bacterial population to expand and persist in the urinary tract lumen. In the laboratory, a single *E. coli* strain can form qualitatively and quantitatively distinctive biofilms with different media composition, temperature, and flow conditions [16, 17].

*Escherichia coli* is the predominant bacterial species associated with asymptomatic bacteriuria, uncomplicated UTI, CAASB, and CAUTI [18]. Unlike enteric pathotypes, there is no definitive genetic signature of a “uropathogenic” *E. coli* strain. Studies have identified *E. coli* characteristics that are common in the setting of infection but none that are definitive of a uropathogenic pathotype, consistent with the view that the uropathogenic potential of *E. coli* is multifactorial in nature [19]. A thorough accounting of infection-associated genes – virulence factors (VFs) – has not been conducted in catheterized patients, though some VFs have been well-documented in CAASB-associated *E. coli* isolates. The VFs identified to date are functionally diverse and may contribute to pathogenic potential differently in catheterized, compared to non-catheterized, patients. Rather than achieving a strict monogenic definition for uropathogenic *E. coli*, the data to date support a probabilistic and combinatorial relationship between virulence determinants and disease.

In the present study, we compared *E. coli* isolates from patients with CAUTI and CAASB. To minimize geographic and temporal variation, these strains were obtained at a single location over a common period of time. Comparisons were based on whole genome sequencing analyses and quantitative biofilm phenotyping using a simulated catheter-biofilm system.

Comparative genomic analyses were used in conjunction with network community analysis to identify gene combinations associated with infection and catheter biofilm formation. CAUTI strains were associated with sequence type 131, a lineage with high antibiotic resistance and distinctive virulence genes. Multiple gene communities were associated with high catheter biofilm formation. The genes most highly associated with biofilm formation encode the ferric citrate uptake system (Fec), which exhibited a causative relationship with catheter biofilm in reporter construct and targeted deletion mutagenesis experiments. Finally, we used a competitive catheter biofilm assay to identify CAASB strains capable of preventing colonization by CAUTI-associated, ST131 isolates.

## Results

### E. coli isolates

To compare *E. coli* strains associated with catheter-associated urinary tract infections (CAUTI) or catheter-associated asymptomatic bacteriuria (CAASB) in hospitalized patients, we identified 62 catheter-associated isolates (18.4%) from a previously described collection of 337 urinary isolates from patients at Barnes-Jewish Hospital/Washington University Medical Center between August 1^st^, 2009 and July 31, 2010 [5, 6]. Of these 62 isolates, 12 met symptom criteria for CAUTI (concurrent fever, T > 38℃) and 16 met criteria for CAASB (lack of fever or other clinical symptoms). As an additional comparator group, 13 *E. coli* isolates corresponding to asymptomatic rectal colonization were collected by rectal swabs from healthy, adult volunteers at Barnes-Jewish Hospital/Washington University Medical Center from 2014-2015, designated as rectal colonizers (RC) (Table 1). In total, 44 *E. coli* isolates were collected for this study, with each isolate from a unique catheterized patient or healthy volunteer. CAUTI and CAASB subjects were of similar age and BMI but exhibited a significant sex difference (*P* = 0.0093). Bacteriuric inpatient subjects were older than non-hospitalized asymptomatic RC subjects (*P* = 0.0309), typical of inpatients in the United States [20]. Moreover, *E. coli* strains isolated from urinary bacteriuria were identified with higher trimethoprim/sulfamethoxazole (TMP/SX) and quinolone resistance in clinical laboratory tests (P < 0.01), consistent with multidrug resistance facilitating urinary colonization in catheterized patients.

**Table 1.** Comparison of study cohort demographics.

| Variable<sup>a</sup> | Clinical cohort<sup>b</sup> |  |  | Two-tailed Fisher's exact test (*P*)<sup>c</sup> |  |
| --- | --- | --- | --- | --- | --- |
|  | CAUTI<br>(n = 12) | CAASB<br>(n = 16) | RC<br>(n = 13) | (CAUTI + CAASB) vs RC | CAUTI vs CAASB |
| Sex (female) | 2 (17%) | 11 (69%) | 7 (54%) | 0.7442 | 0.0093 |
| Age in years (>= 65) | 6 (50%) | 7 (44%) | 1 (8%) | 0.0309 | 1.0000 |
| Body mass index in kg/m<sup>2</sup> (>= 25) | 8 (67%) | 6 (38%) | 8 (62%) | 0.5240 | 0.7022 |
| TMP/SMX<sup>R</sup> | 6 (50%) | 9 (56%) | 0 (0%) | 0.0011 | 1.0000 |
| Fluoroquinolone<sup>R</sup> | 11 (92%) | 9 (56%) | 0 (0%) | 0.0001 | 0.0882 |

<sup>a</sup> TMP/SMX<sup>R</sup>, resistance to trimethoprim/sulfamethoxazole antibiotic. Fluoroquinolone<sup>R</sup>, resistance to fluoroquinolone antibiotics.
<sup>b</sup> CAUTI: catheter-associated urinary tract infection. CAASB: catheter-associated asymptomatic bacteriuria. RC: rectal colonizer.
<sup>c</sup> *P* <= 0.05 is considered statistically significant.

### Phylogenomic analysis

We characterized the genome composition of all 41 *E. coli* isolates using a whole-genome sequencing approach. Of the 15,993 genes identified in the pan-genome of these isolates, 2458 were identified in 100% of isolates, 3014 in 98% (40/41) of isolates, and over 4030 in 50% or fewer isolates. Each isolate was recognized as genetically distinct by comparing their genome differences, without clonal pairs. A maximum-likelihood tree (Figure 1) established based on variable gene content divided isolates into four main clades corresponding to the canonical *E. coli* phylotypes B2, F, D and a combination of A, B1, and E [21]. Nearly all CAUTI isolates (11/12) belonged to phylotype B2 as is typical of extraintestinal *E. coli* [2, 3]. CAASB strains were more broadly distributed among all detected phylotypes with the exception of phylotype E. RC isolates were distributed among phylotypes A, B2, and F. Of note, CAUTI strains clustered at the extreme of the phylogenetic distribution, corresponding to a sub-clade within phylotype B2 (designated as B2a), that significantly separates from non-CAUTI isolates (10/15 vs. 1/13, *P* = 0.0021, two-tailed Fisher’s exact test). The B2a subclade consisted of 14 ST131 strains [22, 23], while the non-B2a isolates (designated as B2b) were more diverse and consisted of eight sequence types (ST 12, 73, 95, 127, 141, 144, 357, 538. Sparse principal components analysis (sPCA) [24, 25] of B2 strain genome composition similarly distinguished B2a from B2b strains, with clear separation on the PC1 (31%) in the score plot (**Supplementary Fig. S1a&b**). Classification of these B2 subclades by logistic regression using PC1 values yielded a prediction accuracy of 1.0 (**Supplementary Fig. S1c**, SD = 0) and an AUC of 1.0 (**Supplementary Fig. S1d**, SD = 0) in five-fold cross validation. These analyses identify robust differences in *E. coli* gene composition in the CAUTI-associated B2a subclade.

**Fig. 1.**
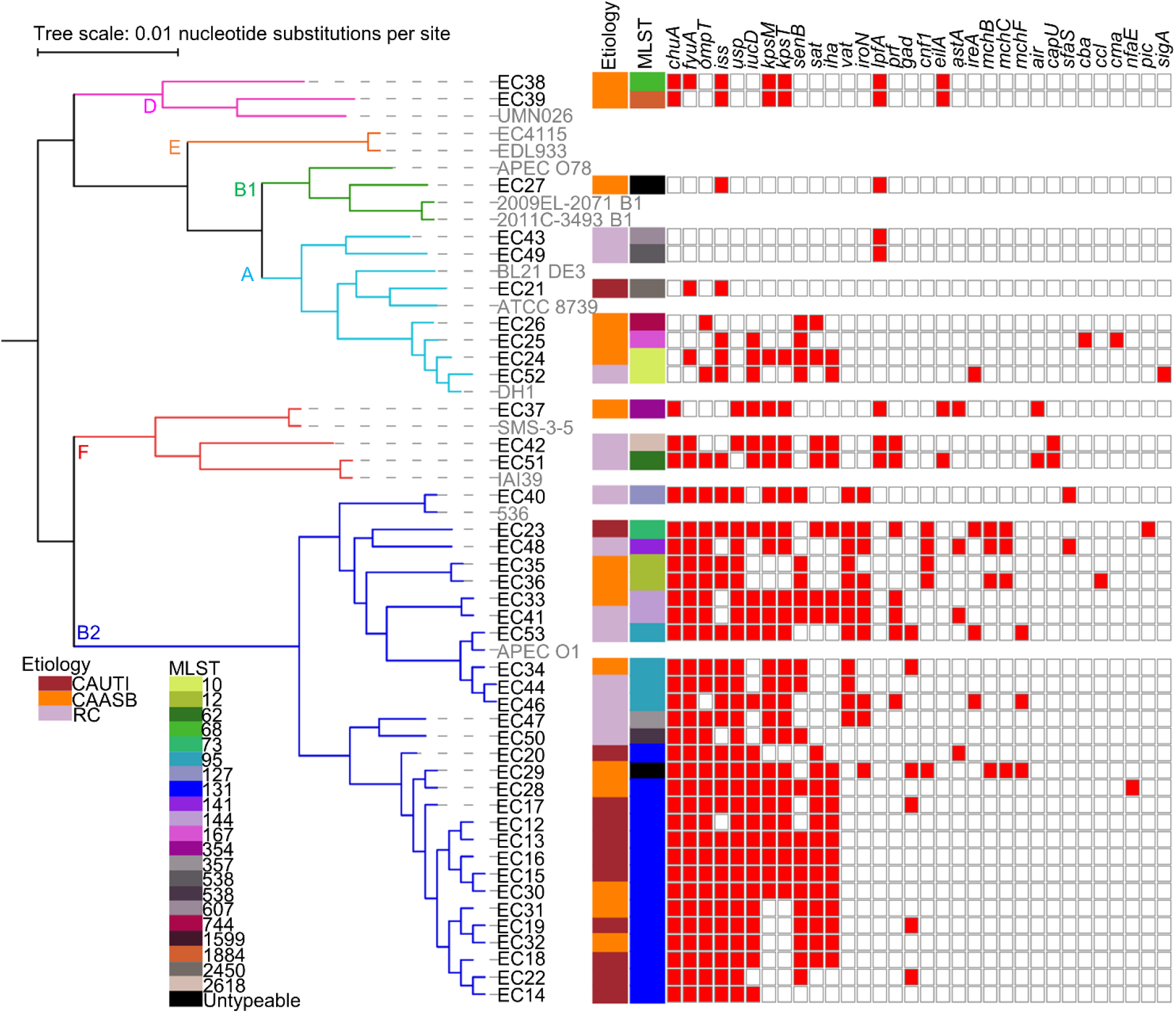
Phylogenetic distribution of 41 clinical *E. coli* isolates. Core-genome alignment was constructed into a maximum likelihood tree with raxML and viewed in iTOL, with six phylotypes identified. *In silico* multilocus sequence types (MLST) were identified using BLASTN to the *E. coli* MLST database, with 21 STs identified. Virulence factors (VFs) were annotated in the *E. coli* draft genomes using VirulenceFinder v1.5 and blastp to previously described genes, with 32 VFs identified. The strain names of *E. coli* sequenced in this study were in black and reference *E. coli* strains were in gray.

### Antibiotic resistance

The ST131 isolates that dominate B2a are a globally emergent extraintestinal pathogenic *E. coli* lineage associated with multidrug resistance, most notably to the fluoroquinolone class of antibiotics [22, 23]. To determine whether increased antibiotic resistance is associated with B2a, we assessed antibiotic resistance gene (ARG) content and phenotypic resistance reported by the clinical laboratory. ARGs against aminoglycosides, beta-lactams, amphenicols, TMP/SX, macrolides/lincosamides/streptogramins (MLS), quinolones, and tetracyclines were identified from the genome assembly (**Supplementary Table S1**). Both phenotypic and genotypic fluoroquinolone resistance were more common in B2a than B2b strains (Table 2, *P* = 0.0001). Specific SNPs previously associated with fluoroquinolone resistance in H30 subclones of ST131 strains (*gyrA* D87N, S83L; *parC* E84V, S80I; *parE* I529L) were nearly ubiquitous in B2a isolates (Table 2, *P* = 0.0001). Moreover, B2a group strains exhibited higher frequencies of ARGs associated with TMP/SX, beta-lactams, and aminoglycosides (**Supplementary Table S1**, *P* < 0.03). Together, the high frequency of resistance genes, particularly those related to fluoroquinolones, is consistent with previously described ST131 isolates [22, 23].

**Table 2.**
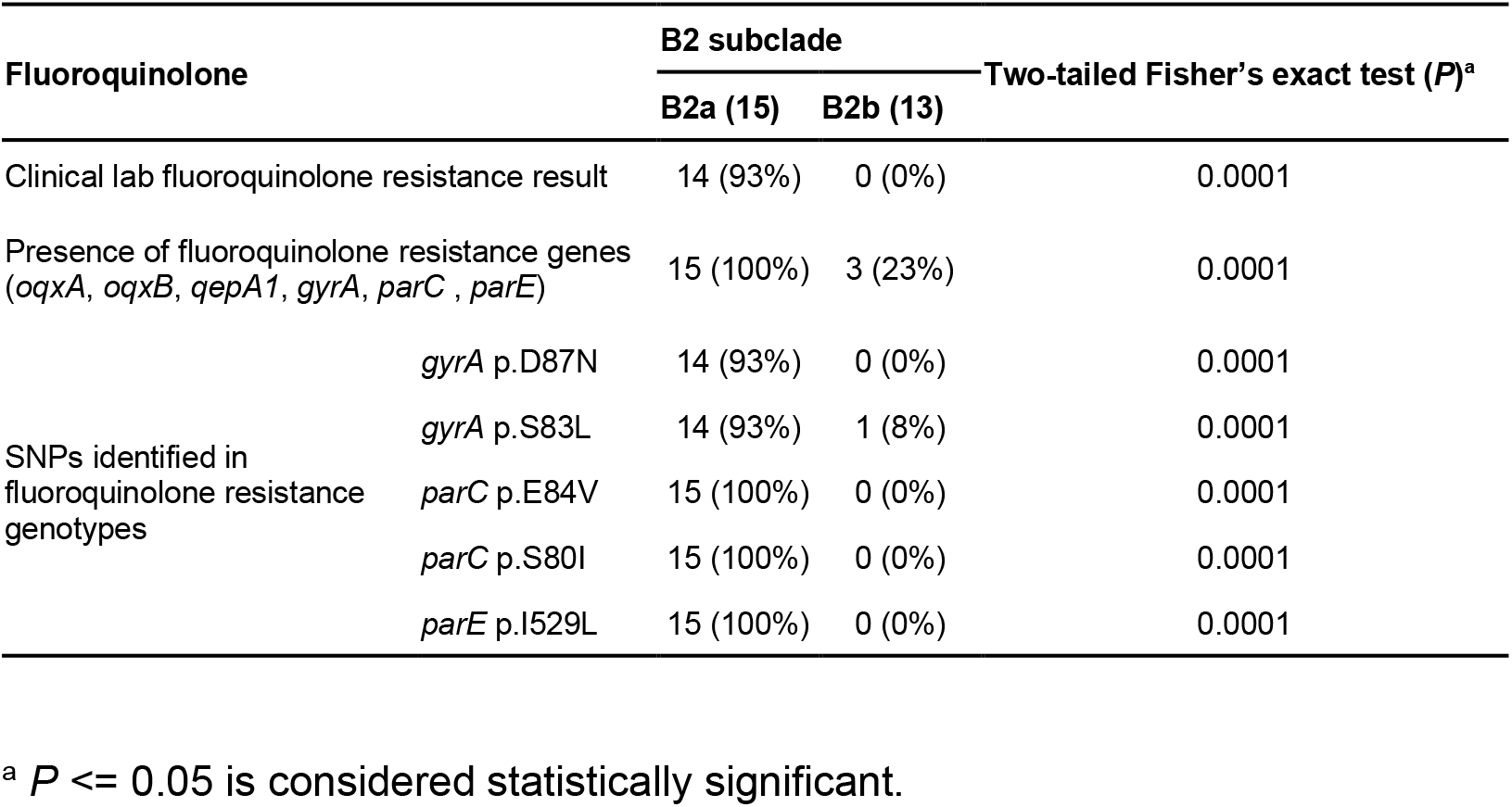
Assessment of fluoroquinolone resistance in phylotype B2 *E. coli* strains.

### Virulence factor content of CAUTI strains

In this cohort, we considered that gene combinations carried by B2a isolates may enhance their pathogenic potential in catheterized patients. Virulence factor genes (VFs) associated with pathogenic gains of function were identified from a list derived from bacterial pathogenesis literatures [26]. We identified 32 such VFs in our isolates (Fig. 1). The number of VFs per isolate, previously known as the “virulence score” [25], did not distinguish (*P* = 0.147) CAUTI (9.5 ± 3.8), CAASB (8.8 ± 3.6), or RC (9.5 ± 4.3) isolates. B2 strains exhibited higher virulence scores than non-B2 strains (10.7 ± 2.7 vs. 5.8 ± 4.0, *P* = 0.002), with a non-significant trend toward a lower score in B2a than B2b (10.0 ± 2.4 vs. 11.5 ± 3.0, *P* = 0.138) (**Supplementary Fig. S2**).

We next considered that VFs affect pathogenic potential through additive or synergistic combinations with other VFs [26]. To assess this, we used network community detection to identify co-associations between the 32 VFs. We performed modularity-based community detection using the Louvain method on a weighted network of positive correlations to assess correlations between the 26 VFs that were present more than once (> 2.4%) among our isolates [27, 28]. Three gene communities were resolved, as visualized by the force-directed network layout (Fig. 2a) and the corresponding correlation matrix (Fig. 2b). Each community was composed of functionally diverse VFs, with iron acquisition systems and toxins prominent in communities 1 and 2, and adhesins in all three. These communities are consistent with the evolutionary accumulation of VFs with additive or synergistic functions.

**Fig. 2.**
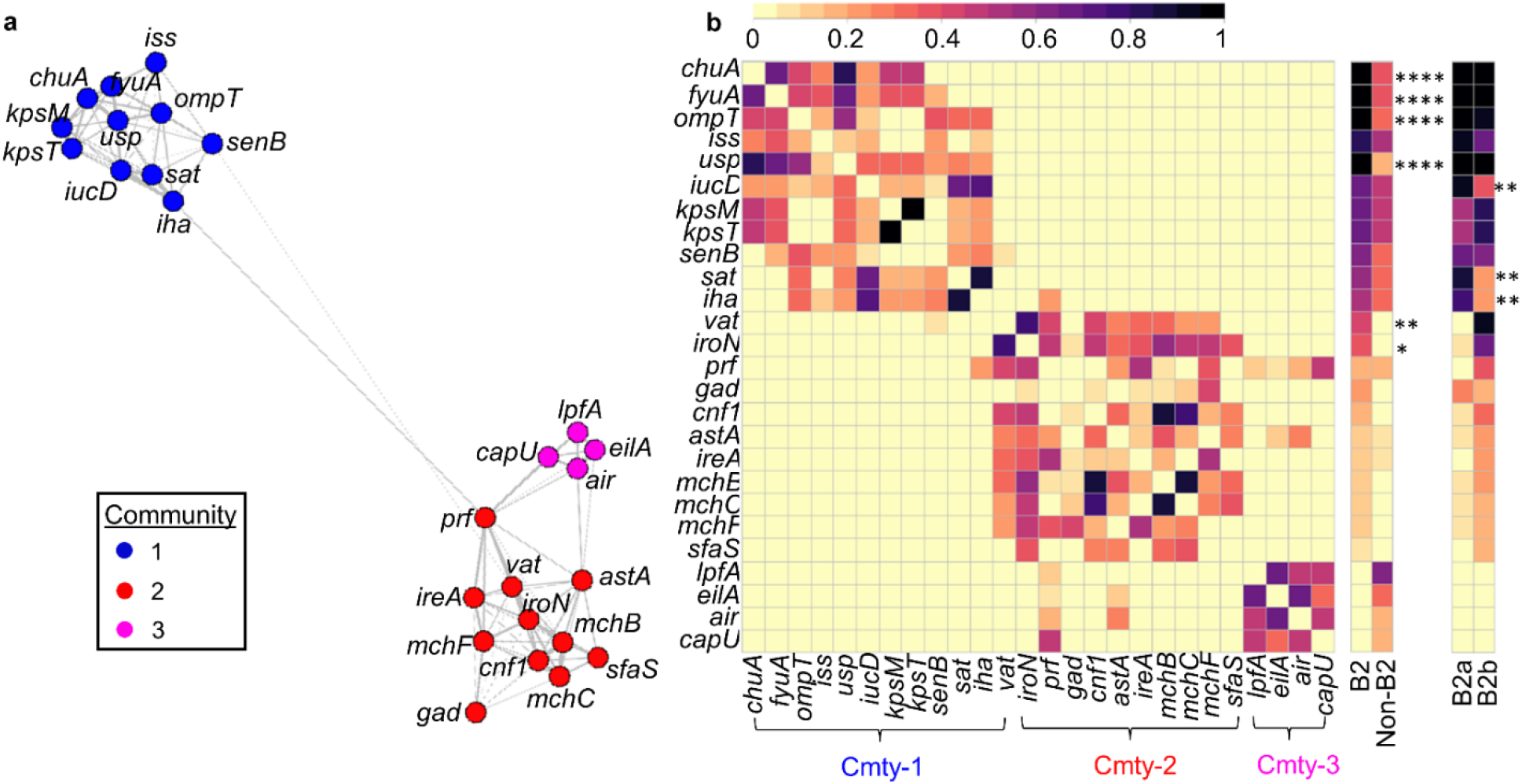
Network analysis of *E. coli* virulence factors (VFs). **(a)** A force-directed network layout illustrated co-associations and three VF communities among 26 VFs. Each node represented a VF. Each connecting line (edge) represented a positive association between 2 VFs that satisfied the significance threshold (0.4% *P*-value threshold, one-tailed on the right, Fisher’s exact test). Edge lengths were determined by the level of correlation between connected VFs. Nodes were colored by community assignment. **(b)** Three VF communities were discernible in the correlation matrix heatmap depicting statistically significant positive associations between 26 VFs. Presence frequency comparisons of each gene between different genetic groups, B2 vs non-B2 and B2a vs B2b, were displayed in heatmaps to the right of the correlation matrix. Cmty: community. By two-tailed Fisher’s exact with *P* <= 0.05 considered statistically significant. *: *P* <= 0.05. **: *P* < 0.01. ***: *P* < 0.001. ****: *P* < 0.0001.

Community 1 and 2 VFs were more common in phylotype B2 strains, while community 3 VFs were exclusively associated with non-B2 strains. Of note, VFs *fyuA*, *chuA*, *ompT*, and *usp* [29–31] were nearly ubiquitous and more common in B2 strains (*P* < 0.00002). Notably, reciprocal relationships between VFs in groups B2a and B2b were evident. Community-1 VFs *iucD, sat*, and *iha* [32–34] were more common in B2a than in B2b (*P* < 0.008). Together, these results are consistent with one set of VFs that raise the pathogenic potential of B2 strains and another set of VFs raising the pathogenic potential of group B2a strains in catheterized patients.

### Biofilm formation by CAUTI, CAASB, and rectal isolates

Catheter biofilm formation is regarded as an important contributor to *E. coli* pathogenic potential. *E. coli* strains form biofilms that vary in important ways depending on media and available substrates [16, 17]. In this study, we compared biofilm formation between *E. coli* isolates using an *ex vivo*, continuous flow model that simulates the clinical catheter environment in patients (**Supplementary Fig. S3**) [35]. We found that artificial urine medium (AUM) [36, 37] yielded growth kinetics (AUM vs human urine = 7.84 ± 0.19 vs 8.06 ± 0.07, *P* = 0.3, Mann-Whitney test) and biofilm morphology (Fig. 3a&b), that were comparable to filter-sterilized human urine. We used this standardized AUM in the continuous flow model to compare biofilm formation between all 41 *E. coli* isolates.

**Fig. 3.**
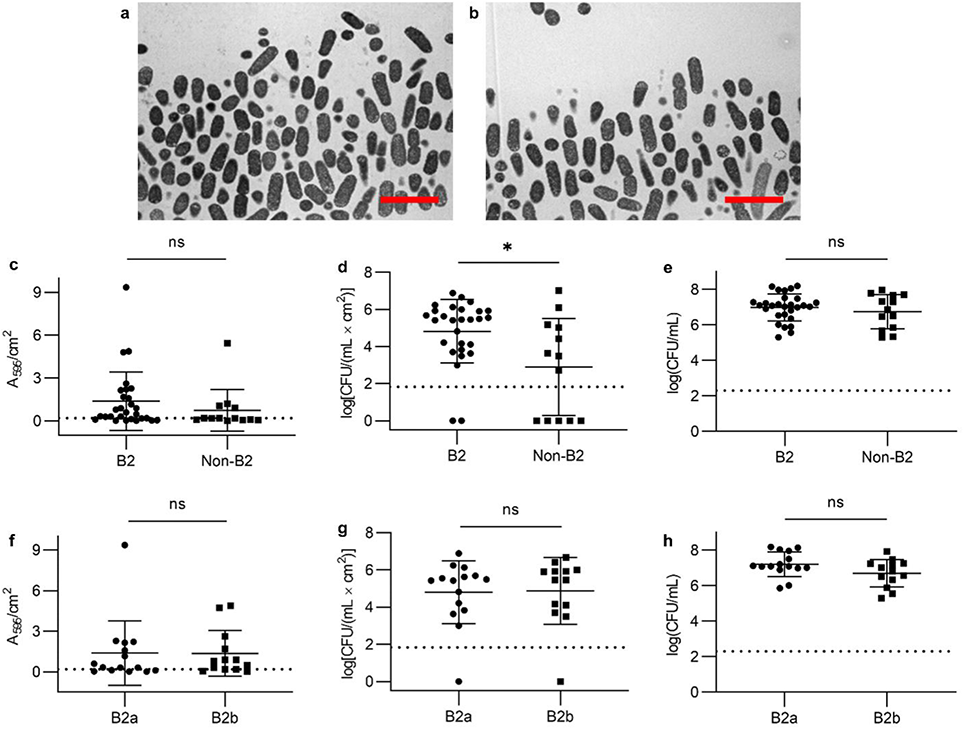
Biofilm formation by CAUTI, CAASB, and rectal isolates. **(a)** Transmission electron microscopy (TEM) image of catheter-biofilm grown in human urine. Scale bar: 4 µm. **(b)** Transmission electron microscopy (TEM) image of catheter-biofilm grown in artificial urine medium (AUM). Scale bar: 4 µm. **(c)** Comparison of biofilm biomass (crystal violet retention) between B2 and non-B2 isolates. Mean with SD plotted for 28 B2 and 13 non-B2 strains. *P* = 0.22. **(d)** Comparison of catheter-adherent CFUs between B2 and non-B2 isolates. Mean with SD plotted for 28 B2 and 13 non-B2 strains. *P* = 0.02. **(e)** Comparison of planktonic CFUs in the voided media between B2 and non-B2 isolates. Mean with SD plotted for 28 B2 and 13 non-B2 strains. *P* = 0.57. **(f)** Comparison of biofilm biomass (crystal violet retention) between B2a and B2b isolates. Mean with SD plotted for 15 B2a and 13 B2b strains. *P* = 0.62. **(g)** Comparison of catheter-adherent CFUs between B2a and B2b isolates. Mean with SD plotted for 15 B2a and 13 B2b strains. *P* = 0.73. **(h)** Comparison of planktonic CFUs in the voided media between B2a and B2b isolates. Mean with SD plotted for 15 B2a and 13 B2b strains. *P* = 0.13. By Mann-Whitney test with *P* <= 0.05 considered as statistically significant. ns: not significant. *: *P* <= 0.05.

Substantial inter-strain variation in catheter biofilm formation was evident, with CV retention (biofilm biomass) ranging from 0.02 to 10.60 A_595_/cm^2^, and adherent CFUs (sessile bacteria in biofilm matrix) from 0 to 10^7.1^ CFU/(mL.cm^2^). Thirty-four of the 41 strains yielded detectable adherent CFUs. Phylotype B2 isolates exhibited significantly higher adherent CFU values (Fig. 3d, *P* = 0.02) with a non-significant trend (Fig. 3c, *P* = 0.22) toward higher CV retention. Neither adherent CFUs nor CV retention was significantly different between groups B2a and B2b (Fig. 3f&g, *P* > 0.6). Differences in planktonic CFUs (planktonic bacteria in voided media) between strains were non-significant in all group-wise comparisons (Fig. 3e&h, *P* > 0.1), regardless of adherent CFU or CV retention values, possibly reflecting bacterial persistence within loosely adherent communities or in turbulent flow. Together, these results are consistent with widespread potential for catheter-biofilm formation among *E. coli*, greater extent of biofilm in phylotype B2, and comparable biofilm formation between CAUTI-associated B2a strains and B2b strains.

### Identification of catheter biofilm-associated genes

To identify catheter biofilm-associated genes, we conducted a comparative genomic analysis of strains with high or low catheter biofilm formation as assessed by CV retention. High and low biofilm isolates were identified within each of the four main clades, B2, F, D, and A + B1(Fig. 1) and designated based on the 95% confidence intervals of their CV retention (biofilm biomass) values, identifying 13 high and 16 low catheter biofilm formers respectively. We next compared genome composition between these groups using sparse partial least squares discriminant analysis (sPLSDA) [38]. In the sPLSDA score plot, high and low biofilm formers were well-resolved along the PC1 axis (Fig. 4a). Classification between high and low biofilm formers by logistic regression using PC1 (9%, **Supplementary Fig. S4a**) values yielded a prediction accuracy of 1.0 (**Supplementary Fig. S4b**, SD = 0) and an AUC of 1.0 (**Supplementary Fig. S4c**, SD = 0) with 5-fold cross validation. Seventy-two genes with the significant PC1 loadings (*P* < 0.05 by two-tailed Fisher’s exact test) were detected, with 46 and 26 genes associated with positive (high-biofilm) and negative (low-biofilm), respectively (Fig. 4b**, Supplementary Table S3**). Of note, the Antigen 43 gene (*flu*) [39] was among the positively associated genes with 44th highest PC1 loading (*P* = 0.04 by two-tailed Fisher’s exact test), providing a confirmatory point of reference to a previously described *E. coli* biofilm-associated gene.

**Fig. 4.**
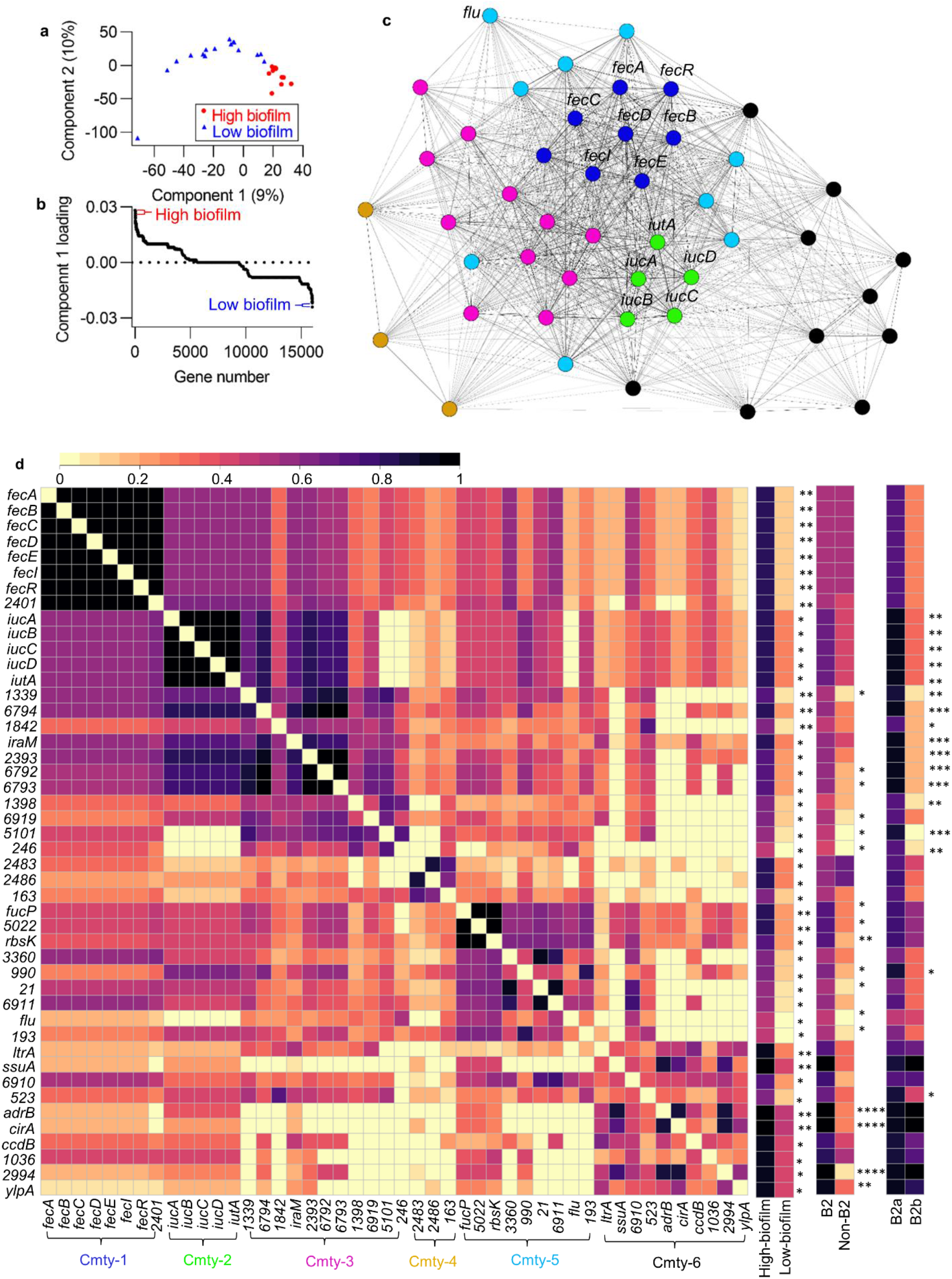
Identification of catheter biofilm-associated genes. **(a)** Score plot of the first two components from sparse partial least squares discriminant analysis (sPLSDA) for displaying group-wise clusterings between high and low biofilm formers. **(b)** Component 1-associated top loadings from sPLSDA identified 72 biofilm-correlated genes, including 46 positive (high-biofilm) and 26 negative (low-biofilm) genes. **(c)** A force-directed network layout illustrated co-associations and three gene communities among 46 biofilm positively associated genes. Each node represented a gene. Each connecting line (edge) represented a positive association between 2 genes that satisfied the significance threshold (5% *P*-value threshold, one-tailed on the right, Fisher’s exact test). Edge lengths were determined by the level of correlation between connected genes. Nodes were colored by community assignment. **(d)** Six gene communities are discernible in the correlation matrix heatmap depicting statistically significant (5% *P*-value threshold, one-tailed on the right, Fisher’s exact test) positive associations between 46 biofilm positively associated genes. Presence frequency comparisons of each gene between different phenotypic and genetic groups, high-biofilm vs low-biofilm, B2 vs non-B2, and B2a vs B2b, were displayed to the right of the correlation matrix. Cmty: community. By two-tailed Fisher’s exact with *P* <= 0.05 considered statistically significant. *: *P* <= 0.05. **: *P* < 0.01. ***: *P* < 0.001. ****: *P* < 0.0001.

To identify co-associations among the 46 genes associated with high catheter biofilms, we performed modularity-based community detection using the Louvain method on a weighted network of positive correlations, as described above for VFs [27, 28]. This analysis resolved six gene communities, as visualized by the force-directed network layout (Figure 4c) and the corresponding correlation matrix (Fig. 4d). Community 1 was composed of the ferric citrate transport (*fecABCDIR*) locus [40, 41] and exhibited robust co-associations and the highest betweenness centrality ranking in the weighted network. The presence of *fec* genes also exhibited the strongest association with high catheter biofilm formation in the sPLSDA analysis (*P* = 0.0025), suggesting a major role in this biofilm phenotype. Community 2 defined the aerobactin siderophore system locus [32] represented by the VF marker gene *iucD*. Relative to communities 1 and 2, communities 3–6 were less robust and are composed of genes associated with more divergent functions.

Together, these results are consistent with a complex polygenic contribution to catheter biofilm production that features a prominent role for iron transport systems (ferric citrate and aerobactin systems, *cir*) and autoaggregation (Antigen 43/*flu*) (**Supplementary Table S3**). The significant positive associations of multiple biofilm genes with B2 and B2a strains (Fig. 4d). suggests that biofilm-associated genes can play a contributing role in the pathogenic potential of CAUTI isolates.

### Fec system activity is associated with increased catheter biofilm formation

The strong association between catheter biofilms and the Fec system raises the possibility that it plays a causative, functional role in enhancing catheter biofilm formation [40, 41]. To address this, we examined the Fec expression profile in a wild type strain and measured catheter-biofilm formation in a *fecA*-deficient mutant. The Fec system imports iron as ferric citrate complexes from the extracellular space to the periplasm through the *TonB*-dependent outer membrane transporter (FecA) [42]. Upon ferric citrate binding, FecA also activates a signaling cascade by interacting with the inner membrane protein FecR which activates the alternative sigma factor FecI. Activated FecI then promotes transcription of other *fec* genes (*fecABCDE*) [40, 41]. To determine whether *fec* gene transcription is associated with *E. coli* biofilm formation, we constructed a FecI-dependent fluorescence reporter strain (EC52::*fecI-RFP*), in the *fec*+ high-biofilm rectal isolate, EC52. Shaking microplate cultures of EC52::*fecI-RFP* exhibited abrupt fluorescence activation at 8 hours when bacteria entered the stationary phase, immediately preceding detectable biofilm formation (Fig. 5a). These observations temporally connect transcriptional activation of *fec* genes to *E. coli* biofilm formation.

**Fig. 5.**
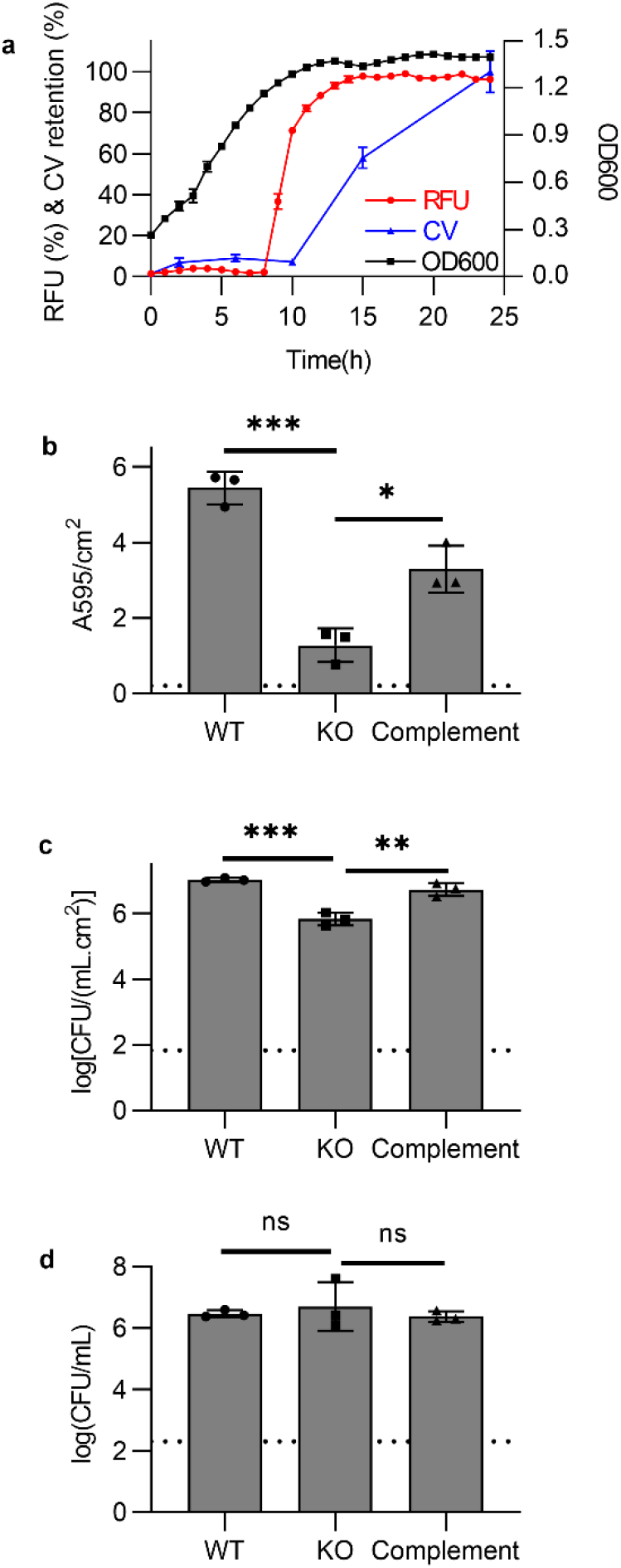
Fec expression is associated with increased catheter biofilm formation. **(a)** Comparison between *fec* expression (RFU), catheter-biofilm formation (CV retention, A595/cm^2^), and bacterial growth (log(CFU/mL)) in the red fluorescence reporter strain, EC52::*fecI*-RFP, measured in a microplate assay. **(b)** Comparison of biofilm biomass (crystal violet retention,) between EC52 (WT), EC52Δ*fecA* (KO), and EC52Δ*fecA*::*fecA* (Complement), measured in the continuous flow model. P < 0.006, by one-way ANOVA with Dunnett’s multiple comparisons test. **(c)** Comparison of catheter-adherent CFUs between EC52 (WT), EC52Δ*fecA* (KO), and EC52Δ*fecA*::*fecA* (Complement), measured in the continuous flow model. P < 0.002, by one-way ANOVA with Dunnett’s multiple comparisons test. **(d)** Comparison of planktonic CFUs in the voided media between EC52 (WT), EC52Δ*fecA* (KO), and EC52Δ*fecA*::*fecA* (Complement), measured in the continuous flow model. *P* = 0.7, by one-way ANOVA multiple comparisons test. Three replicates with mean and SD plotted. *P* <= 0.05 is considered statistically significant. ns: not significant. *: *P* <= 0.05. **: *P* < 0.01. ***: *P* < 0.001.

Next, to determine whether the Fec system affects catheter biofilm formation, we compared catheter biofilm formation by isolate EC52 to its isogenic *fecA* deletion mutant EC52Δ*fecA* [41]. In the catheter biofilm culture system, both CV retention (Fig. 5b, *P* = 0.0001) and sessile bacterial counts (Fig. 5c, *P* = 0.0002) were significantly lower in EC52Δ*fecA* relative to wild type EC52. Genetic complementation of EC52Δ*fecA* with a *fecA* expression plasmid significantly reversed this biofilm formation deficit (Fig. 5b&c, *P* < 0.006). No reduction of planktonic CFUs in the voided media were observed (Fig. 5d, *P* = 0.7). No significant differences in non-biofilm growth were observed between wildtype, mutant and complemented strains (CFU: WT vs KO vs Complement = 7.87 ± 0.058 vs 7.88 ± 0.46 vs 7.84 ± 0.05, *P* = 0.98, one-way ANOVA multiple comparisons test). Together, these data are consistent with a causative, biofilm-specific role for the Fec system in enhancing catheter-associated biofilm formation.

### CAASB strains can inhibit CAUTI colonization

In this study, we found that catheter-biofilm formation was widespread among all *E. coli* isolates, regardless of clinical or genetic groups. Genes associated with biofilm formation are found in CAASB strains and are substantially different from those associated with virulence. This may reflect additional roles for catheter biofilms that do not promote infection. Indeed, the association between biofilm and CAUTI would be weakened if biofilm-forming strains with low pathogenic potential protect the human hosts from infection [43, 44]. A prominent example of such a strain is *E. coli* 83972, a B2 strain isolated from a patient with persistent asymptomatic bacteriuria that has shown efficacy in preventing clinical UTIs following catheter-mediated in of patients [45, 46]. To determine whether CAASB strains can prevent colonization by CAUTI strains, we performed a two-strain competition assay using the continuous flow catheter model. In the continuous flow model, the catheter surface was first pre-colonized by a non-ST131 CAASB biofilm and subsequently challenged with a ST131 CAUTI isolate. The population of each strain was measured using a qPCR method developed in this study, called single-nucleotide polymorphism-selective qPCR (SNPs-qPCR).

Among the 11 non-ST131 CAASB isolates evaluated for competition with ST131 CAUTI isolate EC20, a high-biofilm former, we found that pre-colonization with EC36 (phylotype B2) and EC25 (phylotype A) markedly inhibited both catheter biofilm-adherent (Fig. 6a, *P* < 0.004) and planktonic populations (Fig. 6b, *P* < 0.0003) of EC20 (> 2.5 log inhibition). Pre-colonization with EC36 or EC25 also suppressed the planktonic bacteria growth of the ten CAUTI ST131 isolates (Fig. 6d, *P* < 0.0001, Mann-Whitney test), with isolate EC36 biofilm also significantly inhibiting catheter biofilm colonization (Fig. 6c, *P* = 0.0005). These results demonstrate the ability of a subset of CAASB strains (2/11, 18%) to prevent colonization by CAUTI strains with antibiotic resistance and elevated pathogenic potential [47]. These results raise the possibility that, in a substantial proportion of catheterized patients, *E. coli* catheter biofilm formation may play a protective, not exclusively pathogenic, role.

**Fig. 6.**
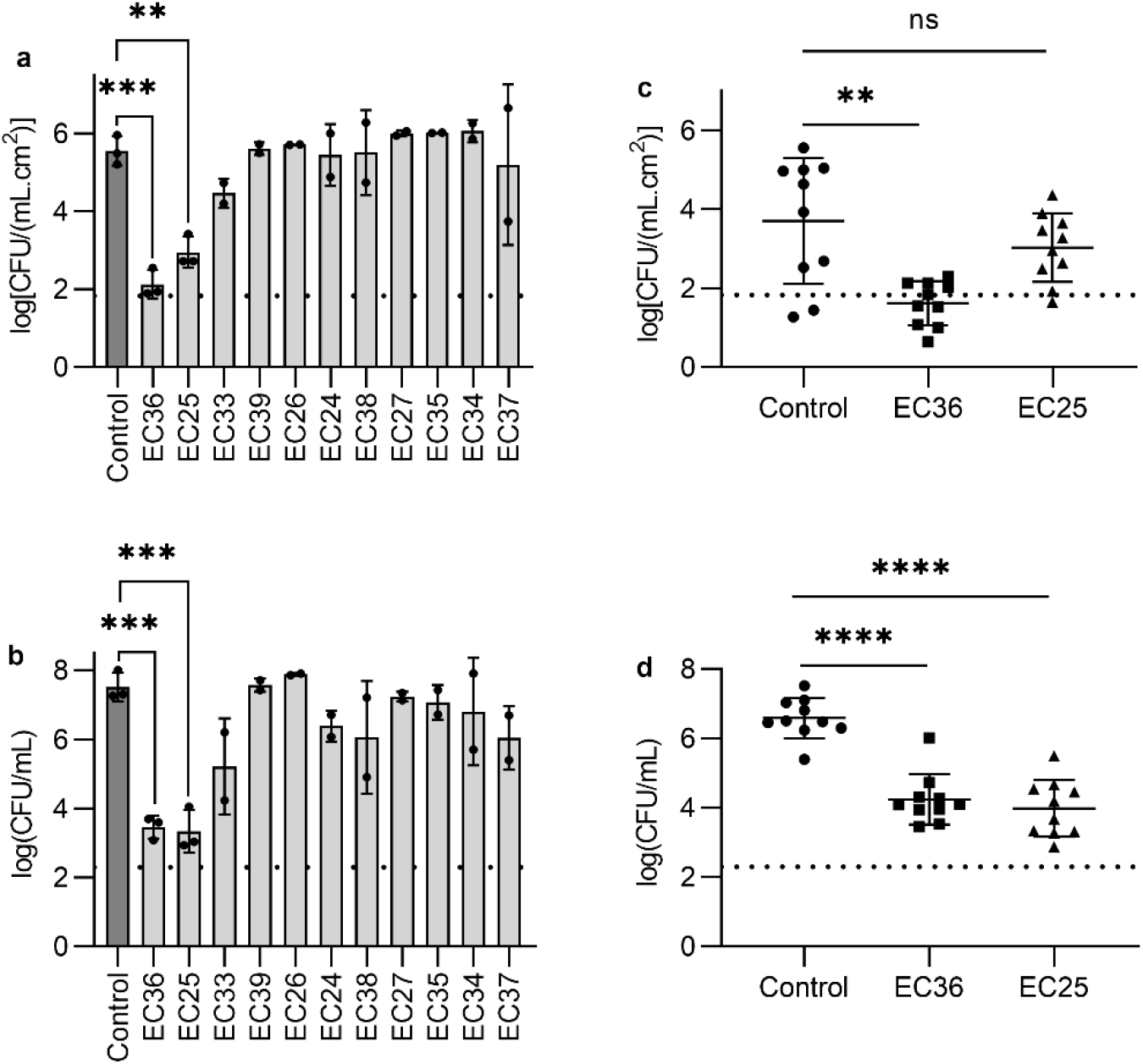
CAASB *E. coli* catheter-biofilms inhibited CAUTI colonization. **(a)** Catheter-adherent CFUs (log[CFU/(mL.cm^2^)]) of strain EC20 in solely biofilm formation test (Control) and in bacterial competition tests with 11 non-ST131 CAASB strains (EC36, EC25, EC33, EC39, EC26, EC24, EC38, EC27, EC35, EC34, EC37). Three replicates with mean and SD plotted for Control, EC36, and EC25, with *P* < 0.004, Two replicates with mean and SD plotted for EC33, EC39, EC26, EC24, EC38, EC27, EC35, EC34 and EC37, with *P* > 0.5. **(b)** Planktonic CFUs (log(CFU/mL)) in the voided media of strain EC20 in solely biofilm formation test (Control) and in bacterial competition tests with 11 non-ST131 CAASB strains (EC36, EC25, EC33, EC39, EC26, EC24, EC38, EC27, EC35, EC34, EC37). Three replicates with mean and SD plotted for Control, EC36, and EC25, with *P* < 0.0003. Two replicates with mean and SD plotted for EC33, EC39, EC26, EC24, EC38, EC27, EC35, EC34 and EC37, with *P* > 0.05. **(c)** Catheter-adherent CFUs (log[CFU/(mL.cm^2^)]) of 10 ST131 CAUTI strains in solely biofilm formation tests (Control) and in bacterial competition tests with two non-ST131 CAASB strains (EC36 and EC25). Test results of 10 ST131 CAUTI strains with mean and SD plotted, with P = 0.0005 and P = 0.3 for EC36 and EC35, respectively. **(d)** Planktonic CFUs (log(CFU/mL)) in the voided media of 10 ST131 CAUTI strains in solely biofilm formation tests (Control) and in bacterial competition tests with two non-ST131 CAASB strains (EC36 and EC25). Test results of 10 ST131 CAUTI strains with mean and SD plotted, with P < 0.0001. By one-way ANOVA with Dunnett’s multiple comparisons test. *P* <= 0.05 is considered statistically significant. ns: not significant. *: *P* <= 0.05. **: *P* < 0.01. ***: *P* < 0.001. ****: *P* < 0.0001.

## Discussion

In this study we identify a genomic lineage within the *E. coli* B2 phylotype that is associated with CAUTI in a hospitalized population. This lineage is dominated by pandemic, multidrug resistant ST131 strains [22, 23], with a distinctive combination of virulence factor genes relative to non-CAUTI-associated isolates. Using a simulated catheter biofilm culture system [35], we found that catheter biofilm formation was not unique to ST131 strains, though catheter biofilms formed in greater extent in the associated B2 phylotype. Comparative genomic analysis revealed a diverse series of catheter biofilm-associated genes, many of which were associated with B2 or ST131 (B2a) isolates [26]. Three iron transport related systems were associated with enhanced biofilm formation and the ferric citrate transport system (Fec), the most prominent of these, was functionally connected to this phenotype through a genetic reporter and targeted deletion mutagenesis. In competitive catheter biofilm experiments, multiple CAASB isolates were capable of diminishing ST131 colonization, suggesting that protection from getting CAUTI may be a relatively frequent consequence of asymptomatic *E. coli* bacteriuria in catheterized patients. The overall results are consistent with enhanced pathogenic potential of *E. coli* ST131 strains in patients with urinary catheters, with biofilm formation playing a contributing, but not determinative role.

Identification of a distinct, infection-associated *E. coli* lineage in a clinical *E. coli* bacteria cohort at this degree of resolution is unusual [19, 25]. This result may reflect the study’s singular focus on catheterized patients, in whom infection may arise through a relatively distinct and uniform pathophysiology in a more homogenous host population. This result may also reflect the relatively recent global proliferation of ST131 strains [23], which may exhibit greater pathogenic potential in catheterized patients relative to strains in previous patient cohorts. The distinguishing characteristic of ST131 strains compared to CAASB strains in this study is not their ability to form catheter biofilms, but rather their carriage of the aerobactin siderophore system, the Sat toxin, and the iha adhesin and the absence of the salmochelin siderophore system and Vat toxin. Strains without this gene profile were more common among CAASB and rectal isolates and some were capable of inhibiting ST131 catheter biofilm colonization.

The ability of some strains to interfere with ST131 catheter colonization raises the possibility that catheter biofilms formed by *E. coli* strains without high-risk virulence gene combinations may benefit patients. Although one such strain that has been extensively studied in this regard, *E. coli* 83972 [45, 46], was collected from an exceptional, non-catheterized patient with three years of bacteriuria, the current study suggests that protective strains may be commonplace in catheterized patients. High antibiotic resistance among ST131 strains [48] raises the possibility that antibiotic treatment may preferentially eliminate protective *E. coli* strains while sparing ST131 strains, paradoxically increasing the likelihood of progression to CAUTI. This scenario further reinforces guideline recommendations to be judicious with antibiotic use [1, 4, 7]. A clinical test distinguishing ST131 from non-ST131 bacteriuria could also aid treatment decisions by helping to differentiate CAUTI and CAASB, a stated area of diagnostic need [48].

The processes that predispose biofilm-bound *E. coli* to cause CAUTI remain unclear but are of diagnostic and therapeutic interest [14]. These processes are presumably complex and include biofilm efflux, host tissue adhesion, immune evasion, and nutrient acquisition [49]. VFs in this study did not appear to equip strains for infection as equally influential components with simple additive effects on pathogenic potential. Neither the number of VFs nor their general functional categories clearly distinguish B2a from B2b strains. B2a and B2b strains are, however, distinguished by the presence of specific VF combinations encoding siderophore, adhesin, and toxin systems. If VFs affect pathogenic potential in catheterized patients, this appears to occur through idiosyncratic functions of specific VFs acting within evolutionarily favored combinations. This is suggested by the presence of VFs in the favored network communities described here (Figure 2) and in previous work [27, 28]. Previously identified VFs are not the only possible contributors to pathogenic potential in ST131 strains. B2a strains in this study carry 224 unique genes that are absent in B2b strains that may also modulate pathogenic potential. Discerning the contributions of these genes would require further study.

The variable genes associated with biofilm formation are substantially different from those associated with CAUTI. In the present study, the ferric citrate uptake system (Fec) was the most prominent of these, with other two iron acquisition systems, the aerobactin siderophore system and the ferric catecholate importer Cir [50], also represented. The deficiency in catheter biofilm formation by a Fec-deficient mutant was surprising, as previous work indicated this deficiency was only discerned in planktonic, siderophore-deficient *E. coli* mutants [51, 52]. These discrepant observations may relate to important differences in iron acquisition and trafficking in the biofilm matrix. The precise nature of these putative differences is unclear. It is possible that the lower iron (III) affinity (relative to siderophores) or lower metabolic cost of citrate as a chelator is an important feature in some biofilm microenvironments. It is also possible that the abundance of host-derived urinary citrate in urine favors bacteria that are able to use this “free” resource [40, 41].

In conclusion, we found that CAUTI in the study population was associated with *E. coli* lineage largely defined by emergent multidrug resistant ST131 strains. The pathogenic potential of these populations was associated with carriage of specific gene networks and high degree of fluoroquinolone resistance, an antibiotic class commonly used to treat UTIs. ST131 strains appeared well-adapted to cause infection in patients with urinary catheters, raising the possibility that these strains arose from co-evolution with catheterized human hosts. The gene networks associated with biofilm formation were largely distinct from the CAUTI-associated gene networks. In addition, catheter biofilm formation was widespread among *E. coli* strains, and some strains in asymptomatic bacteriuria could act to prevent the colonization by CAUTI-associated ST131 strains. These results suggest that strain-specific characteristics of urinary *E. coli* influence CAUTI pathogenesis in patients. Strain-specific testing may thus aid clinical decision making in this population. An improved understanding of how ST131 strains cause infections may suggest future therapeutic strategies for these increasingly antibiotic-resistant bacteria.

## Methods

### Urinary isolate selection

Urinary catheter-associated *E. coli* isolates were identified from the study of bacteriuria (> 5 × 10^4^ CFU/mL) inpatients by Marschall *et al* [5, 6]. This study was approved by the Washington University Institutional Review Board of (WU-IRB). CAUTI was defined as fever (T > 38 °C) with contemporaneous bacteriuria and urinary catheter placement [53]. Documented urinary symptoms (dysuria, lower abdominal pain, flank pain) in the absence of fever were regarded as insufficient for CAUTI diagnosis due to their poor reliability in inpatients, particularly those with urinary catheters [1, 4, 7]. CAASB was defined as bacteriuria in the absence of fever and other documented urinary symptoms.

### Rectal isolate collection

Rectal *E. coli* isolates were collected from healthy, adult volunteers in St. Louis, MO from 2014-2015. This study was approved by the WU-IRB, and all study subjects provided written informed consent. Exclusion criteria included age < 18 years old, pregnancy, current urinary tract infection, previous urogenital surgery, ongoing treatment for urogenital cancer, the use of systemic antibiotics within 30 days of the study visit, or the use of a urinary catheter within 30 days of the study visit. Each study subject used a previously published protocol [54] to procure a self-collected rectal swab (BD Eswab) and submitted it with a study survey. Swabs were processed by the clinical microbiology lab at Barnes-Jewish Hospital to identify a dominant *E. coli* isolate and assess its antibiotic susceptibilities. Fifty-seven subjects were consented, 48 submitted study materials, 41 *E. coli* isolates had matching demographic data, and 13 of those *E. coli* isolates were randomly selected for the current study.

### Human urine collection

Healthy donor urine was collected from adult volunteers as approved by the WU-IRB. Participants provided written informed consent for collection of up to two specimens, at least 1 week apart, for subsequent discovery and validation analysis. Exclusion criteria included recent UTI, antibiotic therapy, pregnancy, or any urogenital diseases. Collected human urine were mixed together, filter-sterilized through 0.22 µm filters, then stored in −80 °C until use. Before an experiment, frozen urine was taken out, thawed on ice, filter-sterilized again and then processed for usage [55].

### Bacterial strains and culture conditions

Isogenic *E. coli* mutant strain was constructed as in-frame deletion in chosen isolate using the Lambda Red recombinase method as described previously [56]. Isogenic mutant complementation and fluorescence reporter construct were accomplished by ectopic expression using transformed plasmids [41]. Unless otherwise specified, cultures were grown from single colonies in LB broth for 12 h at 37 °C before using in the indicated assays.

### Whole-genome sequencing

Bacterial genomic DNA was extracted with the bacteremia kit (Qiagen) from ∼10 colonies of overnight growth. 5 ng of DNA was used as input to create Illumina sequencing libraries using the Nextera kit (Illumina). The samples were pooled and sequenced on an Illumina NextSeq 500 High Output system to obtain 2×150 bp reads. The reads were demultiplexed by barcode and had adapter sequences removed with trimmomatic v.38 and contaminating sequenced removed with deconseq v.4.3 [57]. Processed reads were assembled into draft genomes with SPAdes v3.12.0 (Bankevich). The scaffolds.fasta file from spades was annotated for protein coding sequences on all contigs > 500bp with prokka v1.12 [58]. Additionally, we obtained *E. coli* genomes in the known phylogroups and annotated their protein coding sequences. GFF files from prokka were used as input for roary to create a core-genome alignment with PRANK [59]. The core-genome alignment was constructed into a maximum likelihood tree with raxML and viewed in iTOL [60]. *In silico* multilocus sequence types (MLST) were identified using BLASTN to the *E. coli* MLST database [21]. Previously published virulence factors (VFs) were annotated in the *E. coli* draft genomes using virulencefinder v1.5 and blastp to previously described genes [61]. Antibiotic resistance genes (ARGs) in genomic assemblies were identified by BLAST comparison of protein sequences against the CARD database based on stringent cutoffs (> 95% ID and > 95% overlap with subject sequence) [62]. The genomes analyzed in this report have been deposited to NCBI WGS database under BioProject accession no. PRJNA514354.

### Genomic analysis

The thirty-two virulence factors (VFs) were compared between phenotypic and genetic groups for identifying CAUTI-associated VFs using sparse principal component analysis (sPCA), logistic regression (LR) classification, and network analysis approaches. Biofilm-associated genes were determined by comparative genomic analyses using sparse partial least squares discriminant analysis (sPLSDA), logistic regression (LR) classification, and network analysis approaches [24, 25, 27, 28]. Computational models used in these genomic analyses were configured in Python and R programming languages, mainly by using the scikit-learn module and mixOmics packages, respectively, as well as the Gephi software (http://gephi.org). Because of the high sparsity of genomic metadata, with 15993 genes identified in 41 genome assemblies (15993 >> 41), sparse penalty was enforced in all dimensionality reduction analyses (sPCA and sPLSDA) in the present work to prevent overfitting [63].

### Network analysis

Two network representations for the 26 virulence factors (VFs) and 46 high-biofilm associated genes, connected by co-occurrences across the *E. coli* collection, were defined using statistically significant positive correlations as the edge weights across the networks. Statistical significance between two nodes (genes) was determined by Fisher’s exact test to determine whether they appeared independently, conditional on their observed marginal frequencies among the *E. coli* collection. The 0.4% and 5% *P*-value thresholds (one-tailed on the right) were chosen for the VFs and biofilm-positive genes networks, respectively, to ensure that the obtained gene network in each case was a single connected component. An edge was defined as present between any pair of positively correlated nodes that satisfied the significance threshold, with edge weight equal to the positive correlation coefficient. Communities in this network were detected using the Louvain method by maximizing the modularity function [27, 28]. We selected the obtained 3-community (Resolution = 1.0) and 6-community (Resolution = 1.25) for the network visualizations of 26 VFs and 46 high-biofilm associated genes, respectively, using a force-directed layout generated by the Gephi (http://gephi.org) ForceAtlas2 algorithm and the corresponding correlation matrix.

### Artificial urine medium (AUM)

Artificial urine medium (AUM) (**Supplementary Table S2**) was prepared as an alternative medium of human urine for characterizing biofilm formation. Iron and zinc contents of a previously published AUM recipe [36, 37] was adjusted to reflect that of with human urine specimens measured using inductively coupled plasma-mass spectrometry (ICP-MS), in this study and previous publication [64]. The comparison indicated that the old AUM recipe included more iron (Old AUM vs human urine vs Sieniawska = 5 µM vs 0.86 ± 0.14 µM vs 0.21 µM) and less zinc (Old AUM vs human urine vs Sieniawska = 0 µM vs 9.60 ± 0.89 µM vs 7.0 µM) than pooled human urine. In addition, old AUM recipe without adding 5 µM FeSO_4_.7H_2_O was also measured by ICP-MS, with the results detecting 0.74 ± 0.03 iron and 0.19 ± 0.02 zinc, suggesting that the yeast extract could provide enough iron but not zinc in AUM to mimic human urine composition. The published AUM recipe was therefore modified by removing the 5 µmol/L FeSO_4_.7H_2_O and adding extra 7 µmol/L ZnSO_4_.7H_2_O (**Supplementary Table S2**). All ICP-MS experiments were conducted at the Nano Research Facility (NRF), Department of Energy, Environmental and Chemical Engineering, Washington University in Saint Louis. The ICP-MS quantification was achieved using calibration curve of 1, 5, 10, 50, and 100 µg/L. Nitric acid (Fisher) was added into pooled human urine samples with a final acid concentration of 2% [55].

### Continuous flow biofilm-catheter model system

Biofilms were grown in a continuous flow catheter model, using previously published protocols with appropriate modifications (**Supplementary Fig. S3**) [35]. The components used for assembling continuous flow system included the platinum-cured silicone urinary catheters (Nalgene^TM^ 50), peristaltic pump (Watson Marlow 205U), flexible tubings (Tygon S3^TM^), and plastic connectors (Thermo Scientific). Prior to use, all tubing, connectors and containers were autoclaved. Human urine and AUM were filter-sterilized through 0.22 µm filters. Bacteria from single colonies were grown in LB broth under 37 °C for 12 h, washed with PBS, back-diluted 1:10 into filter-sterilized human urine or AUM, and injected into the catheter installed in the continuous flow system operating under 37 °C. *E. coli* inoculum was statically incubated for two hours to allow the bacterial attachment to catheter surface. Subsequently, fresh medium was pumped through the catheter at the flow rate of 0.5 mL/min, with thirty-minute pre-flush allowed to wash off the loosely adherent bacteria first. Next, at 10 hours of continuous flow incubation, the voided media and catheters were collected for subsequent characterization.

Biofilm was quantified by crystal violet (CV) retention measuring the total biofilm biomass, and CFU assays detecting the sessile bacteria in the biofilm matrix and shedding planktonic bacteria in the void media [35]. Catheter (11 cm) collected from the flow system was washed with PBS and cut into three pieces (3 cm), with two pieces for CV staining and the other one for CFU enumeration. For CV staining, 3 cm catheters were stained in 0.5% CV solution for 10 min, followed by three times deionized water washing to remove the excess dye. The washed catheters were first dried by capillary action on adsorbent paper, and then allowed to air dry at room temperature for overnight. Next, the stained catheters were filled with 33% acetic acid to dissolve bound dye for 10 min. Finally, the acetic acid solutions with dissolved crystal violet were taken out, diluted by 20 times, and then measured at 595 nm using a UV/Vis Spectrophotometer (Beckman Coulter DU-800). The biofilm biomass was quantified as A_595_/cm^2^ by the CV staining method. For sessile bacteria quantification, 3 cm catheter was cut into fragmented pieces, immersed into 3 mL PBS, sonicated (Branson 350) for 10 min, vortexed for 3 min (GeneMate), and proceed for CFU plating to quantify the adhered bacteria detached from the catheter surface. The amount of biofilm bacteria was quantified as CFU/(mL × cm^2^). For planktonic bacteria quantification, voided media were collected and plated for CFU enumeration, quantified as CFU/mL.

### Characterization of biofilm structure

Biofilm grown on urinary catheter surface was processed for structure characterization using transmission electron microscopy (TEM) [65]. For microstructural characterization, 1 cm catheter with biofilm formed on inside surface was fixed in 2% paraformaldehyde/2.5% glutaraldehyde (Polysciences Inc.) in 100 mM sodium cacodylate buffer (pH 7.2) for 1 h at room temperature. Samples were washed in sodium cacodylate buffer and postfixed in 1% osmium tetroxide (Polysciences Inc.) for 1 h. Samples were then rinsed extensively in deionized water prior to en bloc staining with 1% aqueous uranyl acetate (Ted Pella Inc.) for 1 h. Following several rinses in deionized water, samples were dehydrated in a graded series of ethanol and embedded in Eponate 12 resin (Ted Pella Inc.). Sections of 95 nm were cut with a Leica Ultracut UCT ultramicrotome (Leica Microsystems Inc.), stained with uranyl acetate and lead citrate, and viewed on a JEOL 1200 EX transmission electron microscope (TEM) (JEOL USA Inc.) equipped with an AMT 8 megapixel digital camera and AMT Image Capture Engine V602 software (Advanced Microscopy Techniques)

### fec fluorescence and microplate-biofilm assays

To determine the expression of ferric citrate transport (*fec*) pathway during biofilm formation, a *fec* fluorescence assay was performed using transformed plasmid [41]. The promoter sequence of gene *fecI*, chosen from the gene cluster of *fecABCDIR*, was inserted in one plasmid to manage the expression of red fluorescence mCherry protein. Next, the *fec* fluorescence plasmid was transformed into isolate EC52, a high-biofilm former, for making the fluorescence reporter strain, EC52::*fecI*-RFP (**Supplementary Table S4**). A control group comparator, EC52::RFP, was also constructed using the same plasmid but without incorporated *fec* genes managing the fluorescence protein expression (**Supplementary Table S4**). Primers (**Supplementary Table S5**) used to construct the red fluorescence reporter plasmid, *fecI*-RFP, were designed using OligoEvaluator (Sigma-Aldrich), synthesized using Integrated DNA Technologies, and validated by polymerase chain reaction (PCR) prior to use.

The fluorescence assay was performed using a microplate method to monitor the *fec* expression levels at different stages of bacterial growth in M63 minimum medium over 24 hours. To compare between *fec* expression and biofilm formation, the biofilm growth of reporter strain, EC52::*fecI*-RFP, was also characterized in a microplate test under the same conditions as the fluorescence assay. At 0, 2, 6, 10, 15, and 24 hours of shaking incubation, bacterial cultures were removed from the microplate. To quantify the biofilm formed on the inner surface of microplate wells, the wells were washed using PBS, stained using 0.5% CV solution for 10 min, washed with deionized water three times, dried at room temperature overnight, dissolved using 33% acetic acid for 10 min. Samples were then diluted 20 folds and measured at 595 nm using a UV/Vis Spectrophotometer (Beckman Coulter DU-800). The biofilm formation was quantified as A_595_/cm^2^. All microplate assays were performed using a Tecan microplate reader under 37°C.

### fec mutant biofilm assay

Strain EC52, one high-biofilm former, was chosen for making the isogenic *fec* mutant, as another method to investigate the *fec* pathway affecting catheter-biofilm formation. Gene f*ecA*, encoding the ferric citrate outer membrane receptor, was deleted using Lambda Red recombinase method as described before, creating the isogenic mutant EC52Δ*fecA* (**Supplementary Table S4**) [56]. Mutant strain EC*52ΔfecA* was also genetically complemented using the transformed plasmid, generating EC52Δ*fecA*::*fecA* (**Supplementary Table S4**) as the control-group comparison. The catheter-biofilm formation of EC52, EC*52ΔfecA, and EC52ΔfecA*::*fecA* were measured in the continuous flow system using the same protocol as described above, and compared between each other. Primers (**Supplementary Table S5**) used in the *fecA* deletion and complementation were designed using OligoEvaluator (Sigma-Aldrich), ordered from Integrated DNA Technologies, and validated by following procedures including PCR and gel electrophoresis.

### Bacterial interference test

Bacterial interference between 11 non-ST131 CAASB and 10 ST131 CAUTI isolates (**Supplementary Table S6**) were tested in the continuous flow catheter-biofilm system (**Supplementary Fig. S3b**) with two consecutive flow stages (**Supplementary Fig. S5**) [66]. Urinary catheter was pre-colonized with a low-virulence CAASB biofilm for 10 hours. The CAASB biofilm was then challenged by one high-virulence CAUTI isolate for another 10 hours. At the end of the two consecutive flow stages (10 hours + 10 hours), the voided media and urinary catheters were collected for subsequent quantifications. Using the same protocols as described above, both catheter-bound and planktonic bacteria were first plated for CFU enumeration. Next, bacteria cultures were spun down, with the two-strain pellets collected for determining the populations of CAASB and CAUTI bacteria in the mixed cultures using a real-time PCR (qPCR) method. The qPCR quantification method was established based on employing single-nucleotide polymorphisms (SNPs) for inter-bacteria distinction, with more details described in the “Single-nucleotide polymorphisms (SNPs-qPCR)” section. The populations of CAASB and CAUTI bacteria in both planktonic and biofilm two-strain pellets were quantified and used to determine the efficacy of pre-colonized CAASB biofilm for defending against the colonization by CAUTI isolates. All flow experiments were conducted aseptically under 37 °C using AUM.

### Single-Nucleotide Polymorphisms qPCR (SNPs-qPCR)

The method used for quantifying CAASB and CATUI populations in the two-strain pellets was called the single-nucleotide polymorphisms real-time PCR (SNPs-qPCR) in this study. Housekeeping genes[23] were compared to identify SNPs-containing portions to distinguish between the two strains in a competition assay. As the result, two housekeeping genes *adk* (adenylate kinase) and *gyrB* (DNA gyrase) were identified with strain-specific SNPs that could differentiate between 11 non-ST131 CAASB and 10 ST131 CAUTI isolates. Three SNPs-containing portions in gene *adk* were identified to distinguish CAASB isolates EC24, EC25, EC26, EC27, EC37, EC38, and EC39 from ten CAUTI isolates. Two SNPs-containing portions in another gene, *gyrB*, could differentiate CAASB isolates EC33, EC34, EC35, and EC36 from the ten CAUTI isolates (**Supplementary Table S7**). Primers (**Supplementary Table S5**) amplifying the SNPs-containing portion in each gene were designed using OligoEvaluator (Sigma-Aldrich), ordered from Integrated DNA Technologies, and validated by PCR following gel electrophoresis prior to qPCR tests. Prior to qPCR tests, cellular DNAs were extracted using Wizard Genomic DNA Purification Kit (Promega), and measured by NanoDrop 2000 Spectrophotometer (Thermo Scientific). All qPCR assays were performed on a CFX96 Real-Time System (BIO-RAD). The 20 uL PCR mixture contained 1× iTaq Universal SYBR Green Supermix (BIO-RAD), 0.2 uM of each primer, and 3 ng/uL DNA of each specimen. The standard running conditions consist of a 3 min polymerase activation and DNA denaturation at 95℃, another 10 sec DNA denaturation at 95℃, followed by 40 cycles of a 30 sec annealing at 58.5℃, ended with a melt curve with 5 sec at 65℃ first and 5 sec each at 0.5℃ increase between 65℃ and 95 ℃ [67]. Threshold cycles (*Cq*) were obtained to determine the amount of DNA (ng/uL) for each isolate in the mixed bacteria culture, and the quantification was achieved using the calibration curve [*log(Cq) ∼ log(DNA)*] of 0.09375, 0.1875, 0.75, 1.5, and 3 ng/uL (**Supplementary Table S7**). The DNA percentages was determined based the standard curve and used to calculate the population (CFU) each isolate in two-strain pellets, biofilm or planktonic, collected from the competition experiments.

### Statistical methods

GraphPad Prism 8.0 (GraphPad software) was used to generate graphs and perform statistical analysis in this study. We used Mann-Whitney test for two group comparisons and one-way ANOVA for multigroup comparisons. Dunnett’s tests was used to correct one-way ANOVA multigroup comparisons where appropriate.

## Data availability

The genomes analyzed in this report have been deposited to NCBI WGS database under BioProject accession no. PRJNA514354. Other data that support the findings of this study are available within the paper and its Supplementary Information files or from the corresponding author upon reasonable request.

## Code availability

The computer codes for the analyses in this study are available in Github (https://github.com/QL5001/CAUTI-script).

## Acknowledgments

We thank Wandy Beatty for assistance with transmission electron microscopy (TEM). We thank the Edison Family Center for Genome Sciences and Systems Biology staff, Eric Martin, Brian Koebbe, MariaLynn Crosby, and Jessica Hoisington-López for their assistance in genome sequencing and high-throughput computing. We thank John Wildenthal for critical reading of this manuscript. JPH acknowledges Centers for Disease Control Prevention Epicenters Program Grant (CU54 CK 000162), and National Institutes of Health grants R01DK099534 and R01DK111930. GD acknowledges National Institutes of Health grants U01AI123394 and R01AI155893. PJM acknowledges National Institutes of Health grant R01DK111930. WHM acknowledges the KL2TR002346 – ICTS Institutional Career Development Program and the National Institutes of Health grants UL1TR002345 and 1K08AR076464-01. RFP acknowledges the Monsanto Excellence Fund Graduate Fellowship. The content is solely the responsibility of the authors and does not necessarily represent the official view of the CDC or NIH.

## Author contributions

ZZ and JPH developed the concept, designed the overall study approach and constructed patient cohorts for analysis. ZZ conducted the experiments. WHM conducted the healthy volunteer rectal *E. coli* collection. RFP and GD conducted genome sequencing and alignments. ZZ and GLK conducted reporter construct and targeted mutagenesis. ZZ, PJM, and JPH conducted network analyses. ZZ and JPH analyzed the data and wrote the manuscript.

