## Supplementary information for "Pathogenic potential in catheter-associated *Escherichia coli* is associated with separable biofilm and virulence gene determinants"

**Supplementary figures**

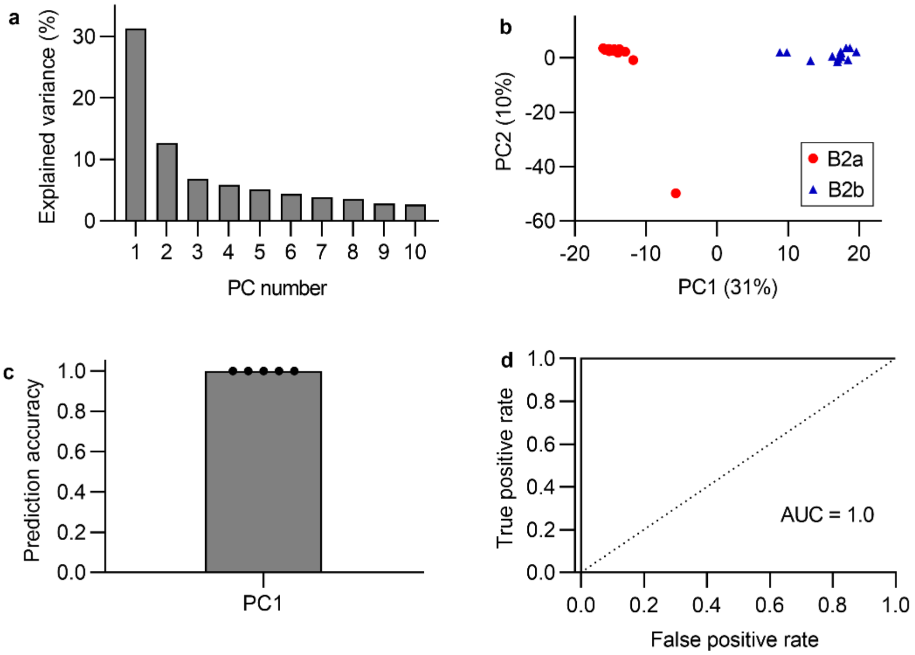


**Supplementary Figure S1. Identification of two genetically different B2 subclades. (a)** Explained variance of the first ten principal components in sparse principal component analysis (sPCA) of phylotype B2 strains gene composition. **(b)** Score plot of the first two principal components for displaying the group-wise clusterings between two B2 subclades, B2a and B2b. **(c)** Logistic regression using sPCA-derived PC1 values for classifying B2a and B2b yielded a prediction accuracy of 1.0 (SD = 0) with 5-fold cross validations. **(d)** Logistic regression using sPCA-derived PC1 values for classifying B2a and B2b yielded an AUC of 1.0 (SD = 0) with 5-fold cross validations.

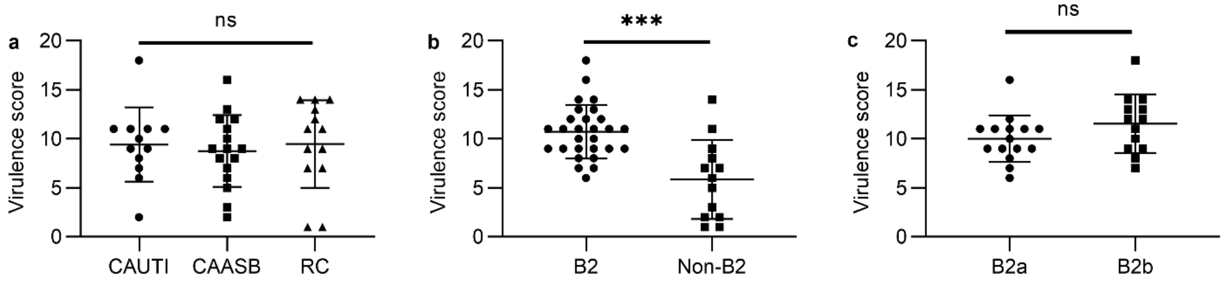


**Supplementary Figure S2. Virulence scores in different phenotypic and genetic groups of *E. coli* strains. (a)** Comparison of virulence scores between phenotypic groups, CAUTI vs CAASB vs RC. Mean with SD plotted for 12 CAUTI, 16 CAASB, and 13 RC strains, respectively. $P$ = 0.147, by one-way ANOVA multiple comparisons test. **(b)** Comparison of virulence scores between phylotypic groups, B2 vs Non-B2. Mean with SD plotted for 28 B2 and 13 non-B2 strains, respectively. $P$ = 0.002, by Mann-Whitney test. **(c)** Comparison of virulence score between subclades within phylotype B2 isolates, B2a vs B2b. Mean with SD plotted for 15 B2a and 13 B2b strains, respectively. $P$ = 0.138, by Mann-Whitney test. $P$ <= 0.05 is considered statistically significant. ns: not significant. ***: $P$ < 0.001.

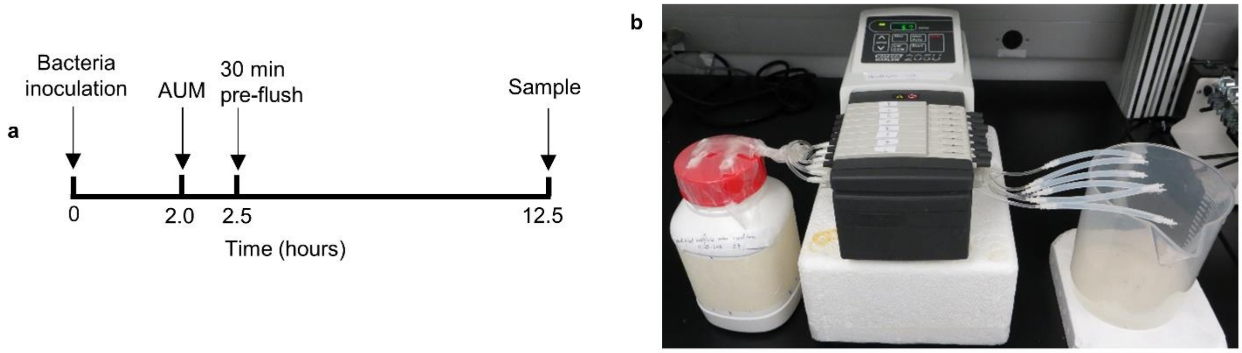


**Supplementary Figure S3. Characterization of catheter-biofilm formation in a continuous flow model. (a)** Experimental timeline of the biofilm formation test. **(b)** Schematic diagram of continuous flow catheter system, including influent, peristaltic pump, urinary catheter, and effluent. AUM: artificial urine medium (AUM)

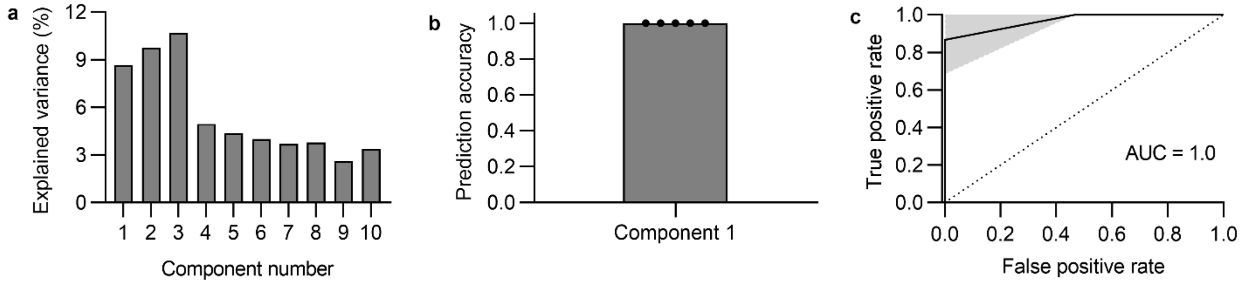


**Supplementary Figure S4. Identification of catheter biofilm-associated genes. (a)** Explained variance of the first ten principal components in sparse partial least squares discriminant analysis (sPLSDA) of gene composition of high and low biofilm *E. coli* strains. **(b)** Logistic regression using sPLSDA-derived PC1 values for classifying high and low biofilm strains yielded a prediction accuracy of 1.0 (SD = 0) with 5-fold cross validations. **(c)** Logistic regression using sPLSDA-derived PC1 values for classifying high and low biofilm strains yielded an AUC of 1.0 (SD = 0) with 5-fold cross validations.

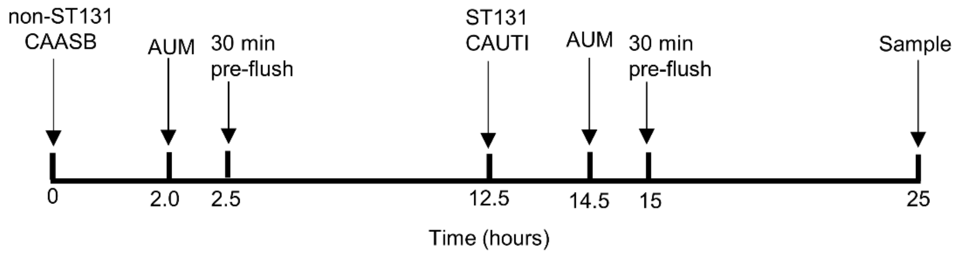


**Supplementary Figure S5.** Experimental timeline of bacterial interference test using continuous flow catheter model.

### Supplementary tables

**Supplementary Table S1.** Carriage of antibiotic resistance genes in phylotype B2 *E. coli* strains.

| **Antibiotic** | **Antibiotic resistance gene** | **B2 subclade** | | **Two-tailed Fisher's exact test ($P$)[a]** |
|---|---|---|---|---|
| | | **B2a (15)** | **B2b (13)** | |
| Aminoglycoside | *aac(3)-IId*, *aac(6')-Ib-cr*, *aac(6')-Ib3*, *aadA1, aadA2*, *aadA5*, *aph(3'')-Ib, aph(3')-Ia*, *aph(6)-Id, strA* | 11 (73%) | 2 (15%) | 0.0032 |
| Beta-lactam | *blaCMY-2*, *blaCMY-7*, *blaLEN16*, *blaOXA-1*, *blaTEM-1B*, *blaTEM-1C*, *ampC* | 15 (100%) | 6 (46%) | 0.0014 |
| Amphenicol | *catA1*, *catA2*, *catB4*, *cmlA1* | 3 (20%) | 0 (0%) | 0.2262 |
| TMP/SMX | *dfrA1*, *dfrA12*, *dfrA15*, *dfrA17*, *dfrA19*, *dfrA5*, *dfrB4*, *sul1*, *sul2*, *sul3* | 9 (60%) | 2 (15%) | 0.0238 |
| MLS | *erm(B)*, *mdf(A)*, *mph(A)* | 15 (100%) | 12 (92%) | 0.4643 |
| Fluoroquinolone | *oqxA*, *oqxB*, *qepA1*, *gyrA*, *parC*, *parE* | 15 (100%) | 3 (23%) | 0.0001 |
| Tetracycline | *tet(A)*, *tet(B)*, *tet(D)*, *tet(M)* | 7 (47%) | 2 (15%) | 0.1145 |

[a] $P$ <= 0.05 is considered statistically significant.

**Supplementary Table S2.** Recipe of the artificial urine medium (AUM)

| Component | Quantify (g) | Concentration (mmol/L) |
|---|---|---|
| Oxoid Peptone L37 | 1 | / |
| Yeast extract | 0.005 | / |
| Lactic acid | 0.1 | 1.1 |
| Citric acid | 0.4 | 2 |
| Sodium bicarbonate | 2.1 | 25 |
| Urea | 10 | 170 |
| Uric acid | 0.07 | 0.4 |
| Creatinine | 0.8 | 7 |
| Calcium chloride.2$H_2O$ | 0.37 | 2.5 |
| Sodium chloride | 5.2 | 90 |
| Zinc sulfate.7 $H_2O$ | 0.00209 | 0.007 |
| Magnesium sulfate.7$H_2O$ | 0.49 | 2 |
| Sodium sulfate.10$H_2O$ | 3.2 | 10 |
| Potassium dihydrogen phosphate | 0.95 | 7 |
| Di-potassium hydrogen phosphate | 1.2 | 7 |
| Ammonium chloride | 1.3 | 25 |
| Distilled water | To 1 Liter | |

**Supplementary Table S3**. Seventy-two biofilm-associated genes.

| **Biofilm level** | **Biofilm-associated genes** |
|---|---|
| High-biofilm (46) | *fecA*, *fecB*, *fecC*, *fecD*, *fecE*, *fecI*, *fecR*, *iucA*, *iucB*, *iucC*, *iucD*, *iutA*, *group_2401*, *group_1339*, *ltrA*, *fucP*, *group_5022*, *group_6794*, *group_1842*, *ssuA*, *group_2483*, *group_2486*, *iraM*, *group_2393*, *group_6792*, *group_6793*, *group_6910*, *rbsK*, *group_1398*, *group_163*, *group_3360*, *group_523*, *group_6919*, *group_990*, *adrB*, *cirA*, *group_21*, *group_6911*, *group_5101*, *ccdB*, *group_1036*, *group_2994*, *ylpA*, *flu*, *group_193*, *group_246* |
| Low-biofilm (26) | *group_7558*, *group_1149*, *group_1713*, *group_228*, *group_312*, *tfaE*, *ascG*, *bglH*, *group_2755*, *tufB*, *gnu*, *group_3170*, *mbtM*, *group_1780*, *hicB*, *rhsB*, *tap*, *yedR*, *group_1225*, *group_1438*, *group_2524*, *group_3689*, *group_405*, *group_5064*, *group_5299*, *group_7629* |

**Supplementary Table S4.** Strains derived for this study to assess *fec* expression and *fec* mutant biofilm formation.

| Strain | Relevant antibiotic resistance[a] | Characteristic | Reference |
|---|---|---|---|
| EC52 | | Wild type *E. coli* rectal colonization isolate, high-biofilm former | (1, 2) |
| EC52::RFP | $Amp^R$ | EC52 ectopically expressing RFP from pGK73 plasmid | This study |
| EC52::*fecI*-RFP | $Amp^R$ | EC52 ectopically expressing RFP from *fecI*-pGK73 plasmid | This study |
| EC52Δ*fecA* | | EC52 with an in-frame deletion of *fecA*, ferric citrate transport deficient | This study |
| EC52Δ*fecA*::*fecA* | $Kan^R$ | EC52Δ*fecA* with an ectopic *fecA* complementation from *fecA*-pKT25 plasmid | This study |

[a] $Amp^R$, resistance to ampicillin antibiotic. $Kan^R$, resistance to kanamycin antibiotic.

**Supplementary Table S5.** Primers used in this study

| Primer Number | Name | Sequence (5'-3') | Description |
|---|---|---|---|
| ZZ#12 | pGK73-Conf FP | TGAACACCATAACCGAAAGTAGT | pGK73 plasmid confirmation forward primer |
| ZZ#13 | pGK73-Conf RP | ACCTTGAAGCGCATGAACT | pGK73 plasmid confirmation reverse primer |
| ZZ#14 | fecI-pGK73-Cons FP | GATCGAGCTCCATCTGATGGAAATGGAAGCCAC | fecI-pGK73 plasmid construct forward primer |
| ZZ#15 | fecI-pGK73-Cons RP | GATCGGATCCCATGCGGAGTGCATCAAAAGTTA | fecI-pGK73 plasmid construct reverse primer |
| ZZ#16 | fecI-pGK73-Conf FP | TGAACACCATAACCGAAAGTAGT | fecI-pGK73 plasmid confirmation forward primer |
| ZZ#17 | fecI-pGK73-Conf RP | TTATTGACATCCTCACTGCCC | fecI-pGK73 plasmid confirmation reverse primer |
| ZZ#18 | fecA-KO FP | TTCTCGTTCGACTCATAGCTGAACACAACAAAAATGATGATGGGGAAGGTATTGTGTAGGCTGGAGCTGC | Red recombinase EC52 fecA knock out forward primer |
| ZZ#19 | fecA-KO RP | CAACATAATCACATTCCAGCTAAAAGCCCGGCAAGCCGGGCGTTAACACAGGTCCATATGAATATCCTCCTTAGTTC | Red recombinase EC52 fecA knock out reverse primer |
| ZZ#20 | fecA-Conf FP | GCGGTACTGGATAAACATTTCAC | EC52 fecA knock out confirmation forward primer |
| ZZ#21 | fecA-Conf RP | CAGGCCTGCAAAAAGAAAAC | EC52 fecA knock out confirmation reverse primer |
| ZZ#22 | fecA-pKT25-Cons FP | GACTAAGCTTGGAAAATAATTCTTATTTCGATTG | fecA-pKT25 plasmid construct forward primer |
| ZZ#23 | fecA-pKT25-Cons RP | GACTGAATTCTCAGAACTTCAACGACCCCTGC | fecA-pKT25 plasmid construct reverse primer |
| ZZ#24 | fecA-pKT25-Conf FP | ATCACATATTCTGCTGACGCA | fecA-pKT25 plasmid confirmation forward primer |
| ZZ#25 | fecA-pKT25-Conf RP | GTTTTACCTGCAGTCCGCTG | fecA-pKT25 plasmid confirmation reverse primer |
| ZZ#26 | adk-24 FP | GATCGTTGACCGTATCGTC | qPCR of adk SNPs-containing region in EC24, EC25, and EC26 forward primer |
| ZZ#27 | adk-24 RP | GTACGGTCTCTTCCTGATCA | qPCR of adk SNPs-containing region in EC24, EC25, and EC26 reverse primer |
| ZZ#28 | adk-27 FP | TGTTGATCGTATCGTCGGT | qPCR of adk SNPs-containing region in EC27 forward primer |
| ZZ#29 | adk-27 RP | AGACGTTTACGCACGGTT | qPCR of adk SNPs-containing region in EC27 reverse primer |
| ZZ#30 | adk-ST131-1 FP | TTGTTGACCGTATCGTAGGC | qPCR of adk SNPs-containing region-1 in ST131-CAUTI isolates forward primer |
| ZZ#31 | adk-ST131-1 RP | TACGGTCTCTTCCTGATCGT | qPCR of adk SNPs-containing region-1 in ST131-CAUTI isolates reverse primer |
| ZZ#32 | gyrB-33 FP | TGACCGAGTTCGAATATGAC | qPCR of gyrB SNPs-containing region in EC33, EC34, EC35, and EC36 forward primer |
| ZZ#33 | gyrB-33 RP | CGTCTTTTTCGGTGGAG | qPCR of gyrB SNPs-containing region in EC33, EC34, EC35, and EC36 reverse primer |
| ZZ#34 | gyrB-ST131 FP | TGACCGAGTTCGAATATGAA | qPCR of gyrB SNPs-containing region in ST131-CAUTI isolates forward primer |
| ZZ#35 | gyrB-ST131 RP | CGTCTTTTTCGGTGGAA | qPCR of gyrB SNPs-containing region in ST131-CAUTI isolates reverse primer |
| ZZ#36 | adk-37 FP | CTCAGGAAGACTGCCGTAAT | qPCR of adk SNPs-containing region in EC37, EC38, and EC39 forward primer |
| ZZ#37 | adk-37 RP | ATGCCCGCTTCTTTCATC | qPCR of adk SNPs-containing region in EC37, EC38, and EC39 reverse primer |
| ZZ#38 | adk-ST131-2 FP | AGGAAGACTGCCGCAAC | qPCR of adk SNPs-containing region-2 in ST131-CAUTI isolates forward primer |
| ZZ#39 | adk-ST131-2 RP | ATGCCCGCTTCTTTCATT | qPCR of adk SNPs-containing region-2 in ST131-CAUTI isolates reverse primer |

**Supplementary Table S6.** Eleven non-ST131 CAASB and Ten ST131 CAUTI *E. coli* isolates selected for bacterial interference tests.

| Group | Strain | Phylotype | Sequence type (ST) |
|---|---|---|---|
| ST131 CAUTI (10) | EC12 | B2 | 131 |
| | EC13 | B2 | 131 |
| | EC14 | B2 | 131 |
| | EC15 | B2 | 131 |
| | EC16 | B2 | 131 |
| | EC17 | B2 | 131 |
| | EC18 | B2 | 131 |
| | EC19 | B2 | 131 |
| | EC20 | B2 | 131 |
| | EC22 | B2 | 131 |
| Non-ST131 CAASB (11) | EC24 | B1 | 10 |
| | EC25 | B1 | 167 |
| | EC26 | B1 | 744 |
| | EC27 | B1 | Untypeable |
| | EC33 | B2 | 144 |
| | EC34 | B2 | 95 |
| | EC35 | B2 | 12 |
| | EC36 | B2 | 12 |
| | EC37 | D | 354 |
| | EC38 | D | 68 |
| | EC39 | D | 1884 |

**Supplementary Table S7.** SNPs-qPCR standard curves for differentially quantifying non-ST131 CAASB and ST131 CAUTI *E. coli* DNAs in mixed bacteria cultures from bacterial interference tests.

| ***log(Cq)* = A*log(DNA)* + B[a]** | | | | | | | | |
|---|---|---|---|---|---|---|---|---|
| ***adk*[b]** | | | ***gyrB*[c]** | | | ***adk*[d]** | | |
| Non-ST131 CAASB | EC24 | y = -0.0841x + 1.2150 | Non-ST131 CAASB | EC33 | y = -0.0775x + 1.1971 | Non-ST131 CAASB | EC37 | y = -0.0808x + 1.2399 |
| | EC25 | y = -0.0796x + 1.2055 | | EC34 | y = -0.0751x + 1.2819 | | EC38 | y = -0.0746x + 1.2921 |
| | EC26 | y = -0.0837x + 1.2036 | | EC35 | y = -0.0904x + 1.1820 | | EC39 | y = -0.0738x + 1.2464 |
| | EC27 | y = -0.0872x + 1.1886 | | EC36 | y = -0.0824x + 1.1836 | ST131 CAUTI | EC12 | y = -0.0751x + 1.2525 |
| ST131 CAUTI | EC12 | y = -0.0803x + 1.2313 | ST131 CAUTI | EC12 | y = -0.0859x + 1.2264 | | EC13 | y = -0.0765x + 1.2707 |
| | EC13 | y = -0.0776x + 1.2508 | | EC13 | y = -0.0795x + 1.2450 | | EC14 | y = -0.0792x + 1.2323 |
| | EC14 | y = -0.0773x + 1.2168 | | EC14 | y = -0.0836x + 1.2132 | | EC15 | y = -0.0793x + 1.2225 |
| | EC15 | y = -0.0777x + 1.2006 | | EC15 | y = -0.0759x + 1.2005 | | EC16 | y = -0.0805x + 1.2153 |
| | EC16 | y = -0.0817x + 1.2068 | | EC16 | y = -0.0706x + 1.2179 | | EC17 | y = -0.0747x + 1.2535 |
| | EC17 | y = -0.0800x + 1.2331 | | EC17 | y = -0.0831x + 1.2230 | | EC18 | y = -0.0688x + 1.2639 |
| | EC18 | y = -0.0749x + 1.2186 | | EC18 | y = -0.0751x + 1.2201 | | EC19 | y = -0.0724x + 1.2664 |
| | EC19 | y = -0.0800x + 1.2442 | | EC19 | y = -0.0785x + 1.2393 | | EC20 | y = -0.0810x + 1.1940 |
| | EC20 | y = -0.0914x + 1.1798 | | EC20 | y = -0.0838x + 1.1845 | | EC22 | y = -0.0681x + 1.2778 |
| | EC22 | y = -0.0747x + 1.2551 | | EC22 | y = -0.0812x + 1.2494 | | | |

[a] Standard curve for quantifying *E. coli* DNAs in mixed bacterial culture from bacterial interference tests. *Cq*: threshold cycles in qPCR. DNA: concentration of DNA, ng/µL.

[b] SNPs identified in one portion of gene *adk* (adenylate kinase) are able to differentiate non-ST131 CAASB strains, EC24, EC25, EC26, and EC27, from ten ST131 CAUTI strains.

[c] SNPs identified in one portion of gene *gyrB* (DNA gyrase) are able to differentiate non-ST131 CAASB strains, EC33, EC34, EC35, and EC36, from ten ST131 CAUTI strains.

[d] SNPs identified in another portion of gene *adk* (adenylate kinase) are able to differentiate non-ST131 CAASB strains, EC37, EC38, EC39, and EC27, from ten ST131 CAUTI strains.
